# Drivers of interlineage variability in mitogenomic evolutionary rates in flatworms (Platyhelminthes) are multifactorial

**DOI:** 10.1101/2022.09.11.507443

**Authors:** Ivan Jakovlić, Hong Zou, Tong Ye, Gui-Tang Wang, Wen-Xiang Li, Dong Zhang

## Abstract

The forces driving interlineage variability in the evolutionary rates (both sequence and architecture) of mitochondrial genomes are often inconsistent and unpredictable. Herein we studied the impacts of multiple variables using 223 flatworm (Platyhelminthes) species and phylogenetic multilevel regression models. We found that: 1. Mitogenomic sequence evolution is faster in parasites associated with the thermally stable environment of endothermic hosts, but the overall impact of thermic habitat is small; 2. Mitogenome sizes are smaller in parasites of endothermic hosts, but the effects are small and inconsistent; 3. Mitogenomic gene order rearrangements (GORR) are positively correlated with mitogenomic size; 4. The expected positive correlation between GORR and sequence evolution is lineage-specific, and non-parasitic species exhibited a strong negative correlation; 5. Longevity has negligible impacts on mitogenomic evolution; 6. Parasitic (Neodermata) flatworm lineages exhibit higher evolutionary rates than non-parasitic lineages; 7. The effective population size has negligible impacts on mitogenomic evolution; 8. Comparatively, parasitism had by far the greatest impact on the mitogenomic evolution, but due to the monophyletic origin of this life-history strategy, alternative hypotheses cannot be rejected. A large number of factors impact the mitogenomic evolution in flatworms, with lineage-specific relative contributions, which sometimes produces incongruent lineage-specific mitogenomic evolution patterns.

## Introduction

Evolutionary rates of mitochondrial genomes (mitogenomes) exhibit remarkable interlineage variability, but the forces driving this variability in the mutation rate appear multifactorial, lineage-specific, and very difficult to predict (Bazin et al. 2006; Montooth and Rand 2008; Nabholz et al. 2008; Galtier et al. 2009; Schaack et al. 2020). Multiple, often conflicting, hypotheses have been put forward to explain the patterns of mtDNA evolution, and different studies often produce conflicting findings (Lanfear et al. 2007; Thomas et al. 2010; Saclier et al. 2018; Jakovlić et al. 2021). We hypothesise that the underlying reason for these contradictory findings and resulting unpredictability is the multifactorial nature of mitogenomic evolution, where a large number of factors impact the evolution with lineage-specific relative contributions.

Mitochondrial genomes of the phylum Platyhelminthes (flatworms) exhibit an amino acid substitution rate more than four times higher than the average substitution rate in the remaining bilaterian taxa (Bernt et al. 2013). A number of other factors also make flatworms an interesting candidate to test different hypotheses related to mitogenomic evolution, such as the associations between sequence evolution and thermic habitat, mitogenome, size, gene order rearrangement rate, longevity, and parasitism (see Table 1, also discussed in more detail in subsequent paragraphs). In previous studies, our team has sequenced and published 20 flatworm mitogenomes, e.g. (Zhang et al. 2018*b*, 2018*c*, 2018*a*), and analysed the evolution of gene order in the Neodermata (Zhang et al. 2019), but to our knowledge, no previous study had attempted a detailed metaanalysis of evolutionary patterns of mitogenomes of Platyhelminthes. In this study, using mitochondrial genomes of over 200 flatworm species, we studied some of the most important variables previously associated with mitogenomic evolution (Table 1) and assessed their relative contributions using phylogenetic multilevel regression models.

**Table 1.**
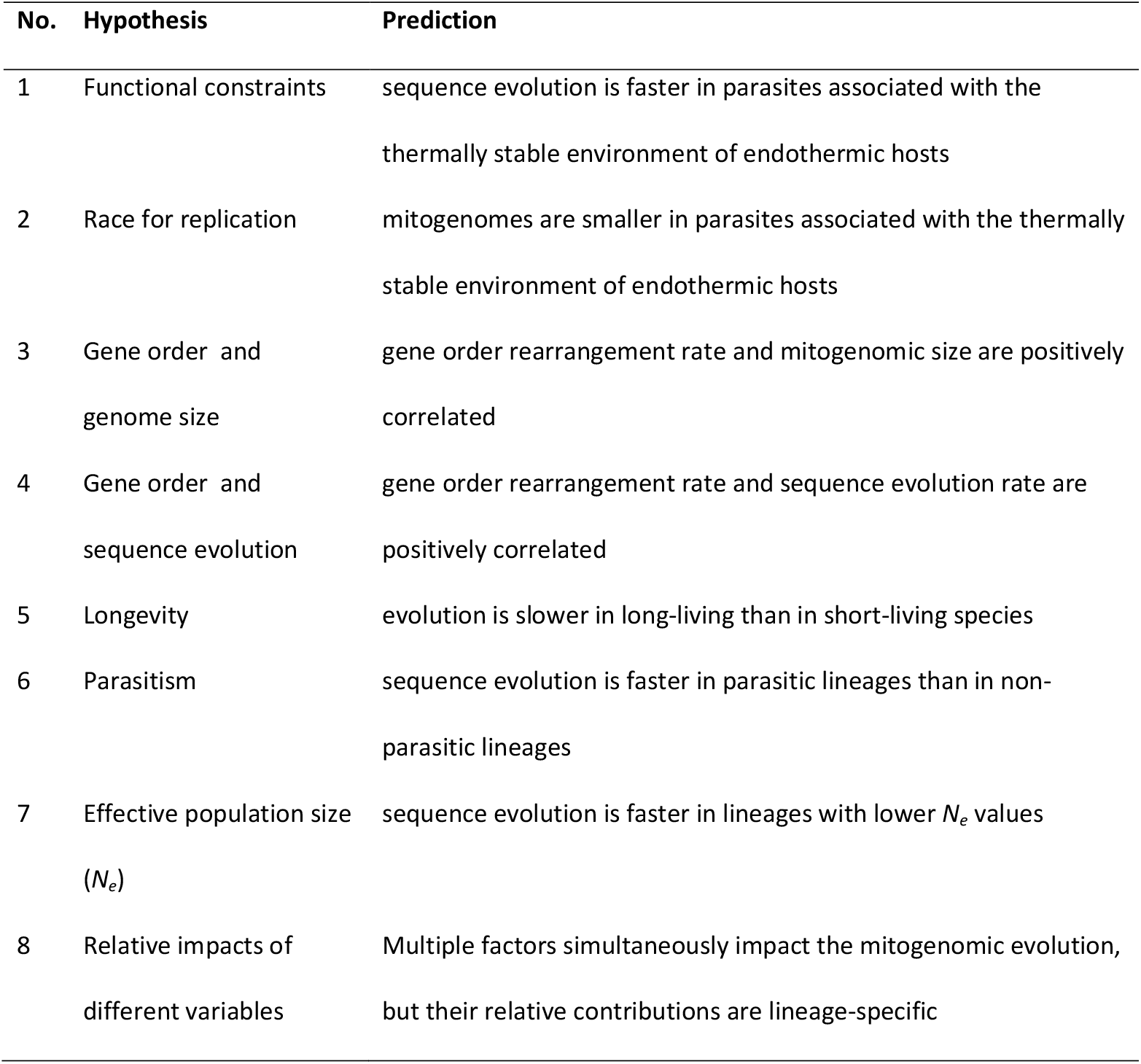
Tested hypotheses and their predictions.

| No. | Hypothesis | Prediction |
| --- | --- | --- |
| 1 | Functional constraints | sequence evolution is faster in parasites associated with the thermally stable environment of endothermic hosts |
| 2 | Race for replication | mitogenomes are smaller in parasites associated with the thermally stable environment of endothermic hosts |
| 3 | Gene order and genome size | gene order rearrangement rate and mitogenomic size are positively correlated |
| 4 | Gene order and sequence evolution | gene order rearrangement rate and sequence evolution rate are positively correlated |
| 5 | Longevity | evolution is slower in long-living than in short-living species |
| 6 | Parasitism | sequence evolution is faster in parasitic lineages than in non-parasitic lineages |
| 7 | Effective population size ($N_e$) | sequence evolution is faster in lineages with lower $N_e$ values |
| 8 | Relative impacts of different variables | Multiple factors simultaneously impact the mitogenomic evolution, but their relative contributions are lineage-specific |

### Hypothesis 1: Functional constraints (thermic habitat and sequence evolution)

Several studies found support for the hypothesis that thermic habitat affects mitochondrial evolution (Melvin and Ballard 2017; Lajbner et al. 2018) and that evolutionary rates differ between endotherms and ectotherms (Rand 1994). Two hypotheses have been put forward to explain this association. The ‘functional constraints’ hypothesis proposes that variations in the thermic environment may restrict physiologically acceptable amino acid substitutions, which implies that thermally stable endotherms should have a higher rate of sequence evolution than thermally variable ectotherms (Thomas and Beckenbach 1989; Rand 1994). The ‘metabolic rate’ hypothesis postulates that endothermic animals have higher metabolic rates than ectotherms, which generates more reactive oxygen species (ROS) and results in an increased DNA mutation rate in endotherms (Martin and Palumbi 1993; Martin 1995). Due to their similar predictions and absence of metabolic rate data for flatworms, we did not have the means necessary to distinguish between the ‘metabolic rate’ and ‘functional constraints’ hypotheses in practice, so herein we nominally refer to both together as the ‘functional constraints’ hypothesis. While all flatworms are themselves ectothermic, some parasitic lineages spend most of their life parasitizing within endothermic hosts, whereas others parasitise ectothermic hosts, so their predominant thermic habitats can be drastically different both in terms of average values and stability. Lagisz et al. proposed that this may also affect mitogenomic evolution, but failed to find evidence in nematodes (Lagisz et al. 2013). Herein, we divided parasitic flatworms (Neodermata) according to the thermic environment of their intermediate and definitive host into ectotherms and endotherms, and tested the hypothesis that sequence evolution is faster in flatworm parasites associated with the thermally stable environment of endothermic hosts.

### Hypothesis 2: Race for replication (thermic habitat and genome size)

The ‘race for replication’ hypothesis proposed that higher metabolic demands associated with endothermy impose stronger purifying selection constraints for small genome size, which would be reflected in reduced mitogenome size and size variability in endotherms (Rand 1993). There is evidence that these patterns are mirrored in some parasitic lineages: mitogenome sizes of parasitic Nematoda species that spend a significant proportion of their life inside endothermic definitive hosts had more compact mitochondrial genomes than those living inside ectothermic hosts (Lagisz et al. 2013). Herein, we used the dataset outlined in the previous section to test the hypothesis that mitogenomes are smaller and less variable in size in parasites associated with the thermally stable environment of endothermic hosts.

### Hypothesis 3: Gene order and genome size

We expected that at least some mitogenomic architecture rearrangements should produce sequence duplications (Boore 2000), which should result in larger overall genome sizes. For example, architecturally hypervariable mitogenomes of some Nematoda lineages also exhibit exceptionally large sizes (Hyman et al. 2011; Zou et al. 2022*b*), and compact genomes are structurally stable in nematodes (Lagisz et al. 2013). Accordingly, we expected that mitogenome size should be positively correlated with the gene order rearrangement rate (GORR). Flatworms exhibit a strong interlineage discontinuity in the evolution of mitogenomic architecture, with lineages with a remarkably conserved architecture interspersed with lineages with highly elevated architecture rearrangement rates (Zhang et al. 2019), and their mitogenome sizes vary from 13 to 27 Kbp, which makes them a suitable model to test this hypothesis. Herein, we used the entire flatworm dataset to test the hypothesis that gene order rearrangement rate and mitogenomic size are positively correlated.

### Hypothesis 4: Gene order and sequence evolution

Gene order rearrangements should be affected by the purifying selection as they may affect the regulation of gene expression (Boore 1999), so we also hypothesised that the above-mentioned positive correlation between GORR and sequence evolution should be a consequence of the overall trend in purifying selection pressures. Indeed, multiple studies found that GORR and mitochondrial sequence evolution are positively correlated (Shao et al. 2003; Hassanin 2006; Xu et al. 2006; Bernt et al. 2013; Zou et al. 2022*b*). Herein, we used the entire flatworm dataset to test the hypothesis that gene order rearrangement rate and sequence evolution are positively correlated.

### Hypothesis 5: Longevity-dependent selection

One of the major variables proposed to explain the variation in the evolutionary rate of mtDNA is linked to the role of mitochondria in the ageing process, so it is named the ‘longevity-dependent selection’ hypothesis. It proposes that long-lived animals may have adapted to an increased lifespan by evolving macromolecular components more resistant to oxidative damages, thus reducing their evolutionary rates (Nabholz et al. 2008; Welch et al. 2008; Galtier et al. 2009). This hypothesis has been thoroughly tested in mammals, and several studies also found support for it in invertebrates (Huang et al. 2008; Moosmann and Behl 2008; Galtier et al. 2009; Thomas et al. 2010), but some studies also observed that there are multiple exceptions from the expected negative relationship between the generation time and molecular clock rate (Rand 1994; Nabholz et al. 2008; Wang and Hekimi 2015; Allio et al. 2017; Saclier et al. 2018). Flatworms exhibit a wide variation in longevity: whereas monogeneans on average have very short life spans of several weeks (Bakke et al. 2007), trematodes have maximum lifespans up to 25 years (Muller and Wakelin 2002), and some non-parasitic planarians are practically somatically immortal (Valenzano et al. 2017). Herein, we used the entire flatworm dataset to test the hypothesis that sequence evolution is slower in mitogenomes of long-living flatworms.

### Hypothesis 6: Parasitism

Multiple studies observed that parasitism may be associated with accelerated mitochondrial evolution rates in Arthropods (Dowton and Austin 1995; Page et al. 1998; Castro et al. 2002; Hassanin 2006; Shao and Barker 2007; Oliveira et al. 2008; Bernt et al. 2013; Jakovlić et al. 2021). Only a few studies reported this phenomenon in non-arthropod lineages (Martin et al. 2000; Bernt et al. 2013), and some studies also found that the association between elevated mitochondrial sequence evolution rates and parasitism is inconsistent (Castro et al. 2002; Bernt et al. 2013), so further studies are needed to test the universality of this phenomenon. While the phylum Platyhelminthes is comprised largely of parasitic lineages, it also comprises a substantial (paraphyletic) basal radiation of non-parasitic lineages. Among more than 220 complete mitochondrial genomic sequences currently (Dec 2021) available in the GenBank database for this taxonomic group, approximately 30 of those belong to non-parasitic species. Unfortunately, the major parasitic radiation, Neodermata, is monophyletic, so flatworms do not represent an ideal lineage to test this hypothesis. Regardless, with this limitation in mind and using algorithms that correct statistical analyses for phylogenetic relatedness, herein we tested the hypothesis that mitogenomic sequence evolution is faster in parasitic than in free-living flatworms.

### Hypothesis 7: The effective population size (Ne)

Populations with small *N*_*e*_ should experience faster rates of evolution than populations with larger *N*_*e*_ because of the increased influence of drift on selection (Ohta 1992), so previous studies proposed that the underlying reason for elevated evolution rates in parasitic lineages may be *N*_*e*_ reductions caused by high speciation rates and frequent founder events during transmissions to new host individuals (Page et al. 1998). Lynch et al. went even further and proposed that mitogenomic evolution in general is driven primarily by nonadaptive forces affected by the *N*_*e*_: random genetic drift and mutation pressure (the nearly neutral theory of molecular evolution) (Lynch et al. 2006). Herein we used the flatworm dataset to test the hypothesis that fast-evolving lineages have lower *N*_*e*_.

### Hypothesis 8: Relative impacts of different variables

We hypothesised that the underlying reason for inconsistency in results among different studies may be attributed to lineage-specific variability in relative contributions of different variables on mitogenomic evolution. More specifically, this hypothesis predicts that different variables will affect the mitogenomic evolution, but with varying magnitudes of effects, so impacts of some variables may be dwarfed by impacts of other variables. Due to the lineage-specific nature of the magnitude of effect, this would explain why mitogenomic evolution was previously described as “practically unpredictable” (Bazin et al. 2006). Herein, we used phylogenetic multiple regression models to test the hypothesis that the magnitude of effects varies strongly among the different tested variables.

## Methods

### Datasets

A total of 223 flatworm mitogenomes were retrieved from the GenBank database (last accessed 25th February 2021). PhyloSuite (Zhang et al. 2020) was used to extract all mitogenomic data, calculate GC skews (for the coding section of the entire mitochondrial plus strand), and generate comparative tables. Among the 223 species, 30 were non-parasitic (comprising the basal flatworm radiation). The Neodermata (parasitic), were divided in two different ways: 1) along the taxonomic lines into the three major parasitic classes: Cestoda, Trematoda and Monogenea (‘phylogenetic dataset’), and 2) according to the definitive host’s thermic type into endotherms (124) and ectotherms (55) (‘thermic dataset’). As many flatworm parasites of endotherms have an ectothermic intermediate host, we first assigned all parasites according to the definitive host type. As the definitive host provides the environment for most of the individual’s growth and reproduction, it has been argued that it should have a greater impact on the mitogenomic evolution of parasites (Lagisz et al. 2013). Following this, we further tested the impact of different host types by subdividing the parasites according to both intermediate and definitive host types. All parasites with ectotherm definitive hosts also had ectotherm intermediate hosts (ecto-ecto), but parasites with endothermic definitive hosts had both endothermic (endo-endo) and ectothermic (endo-ecto) hosts. The thermic environment data were retrieved from a wide range of public resources. Because of the missing data for different variables, some statistical analyses had to be conducted on slightly reduced datasets (Figure 1; more details in Supplementary files S1 and S2). Notably, all datasets comprising the *N*_*e*_ data were strongly reduced due to the basic dataset containing only 77 species. Given the absence of precise longevity data for many flatworm lineages, we first categorised the longevity into six categories, and then coded longevity as a mean of a range in days: 15 (0 – 30 days), 105 (30 – 180 days), 270 (6 months - 1 year), 547 (1 – 2 years), 1278 (2 - 5 years), and 2500 (> 5 years). For a small number of species, we could not find longevity data, so we applied average values for their respective classes (inferred to be the best option on the basis of the general description of the longevity of these lineages and comparison with available data for closely related sequences): 15 for Monogenea, 105 for non-parasitic flatworms, 547 for Cestoda, and 1278 for Trematoda.

**Fig. 1.**
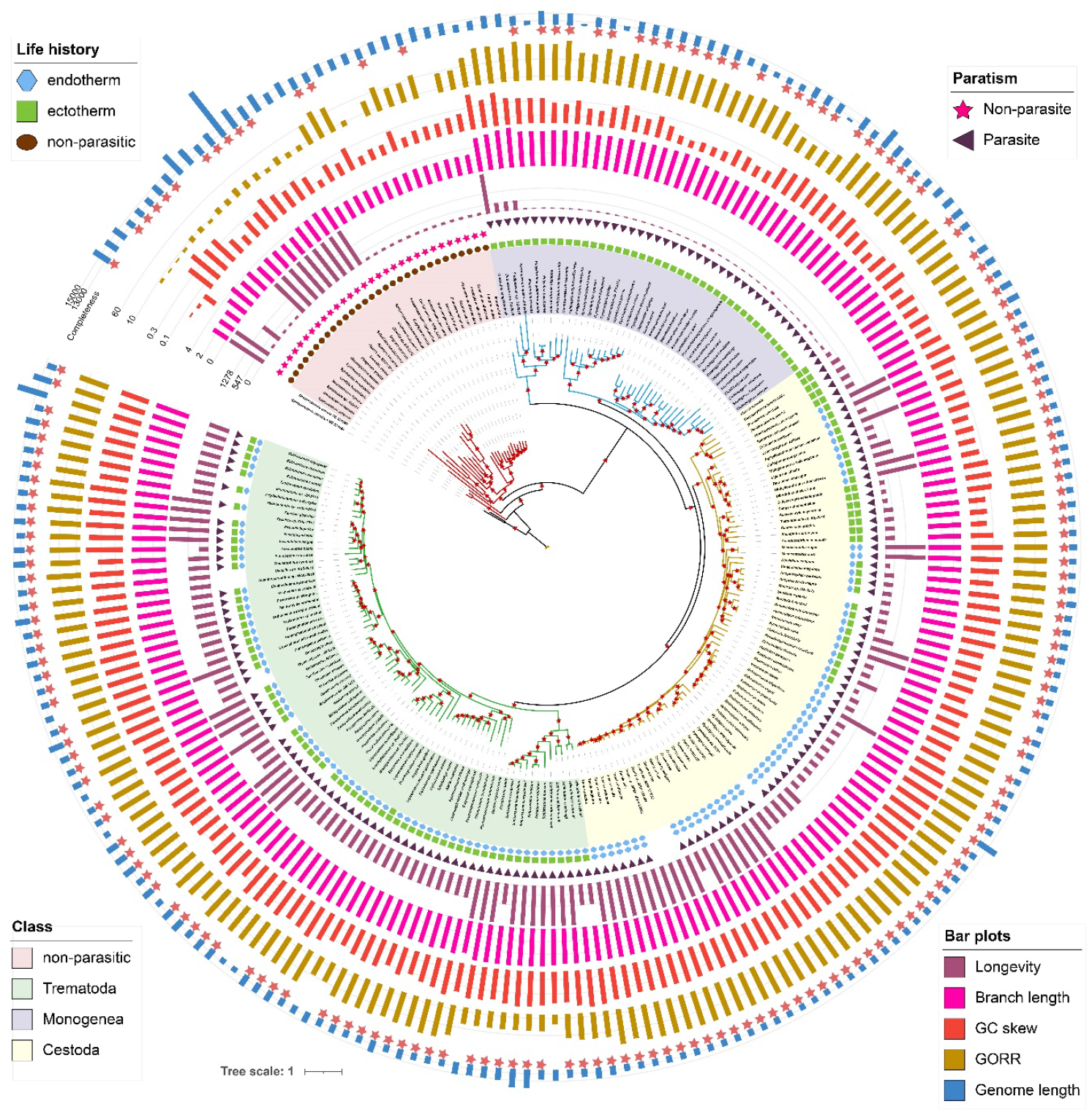
Phylogeny, taxonomy (class), thermic type of definitive host, thermic type of intermediate host, parasitism, longevity, branch length, GC skew, GORR, and genome length of all 223 flatworm species used in the analyses. Parasites are divided into endotherms and ectotherms according to the thermic type of definitive (the inner circle) and intermediate (the outer circle) hosts (the life history legend). The parasitism category shows whether the species is parasitic or free-living. GC skew bars represent the overall skew magnitude on the entire coding section of the plus strand. GORR is the gene order rearrangement rate, where small values indicate highly rearranged mitogenomes. The genome length bar 0 value is set at 12,000 bp, and an absence of a star next to the genome length bar indicates that it was excluded from all analyses that required complete mitogenomes. The star symbols above branches indicate a 100% IQ-TREE ultrafast bootstrap support.

### Phylogenetic analyses

Phylosuite was used to conduct phylogenetic analyses with the help of its plug-in programs on a dataset comprising concatenated sequences (amino acid and nucleotide) of all 12 mitochondrial protein-coding genes (PCGs). We used the methodology described before (Jakovlić et al. 2021) and IQ-TREE (Minh et al. 2020) and PhyloBayes-MPI 1.7a (CAT-GTR model) (Lartillot et al. 2007) (more details in Supplementary files S1 and S2). The molecular evolution rate was defined as the sum of branch lengths (Lanfear et al. 2007; Allio et al. 2017), which were extracted in two steps: the *ape* package was used to root the unrooted tree (Paradis and Schliep 2019) and the patristic method in *adephylo* was used to calculate tip to root distance (Jombart et al. 2010).

### Statistical, GORR, selection pressure, *N*_*e*_, and multivariate regression analyses

To infer the GORR, we first inferred the putative ancestral gene order for Platyhelminthes using MLGO (Hu et al. 2014) with putative sister-group Gastrotricha (Laumer et al. 2019) as the outgroup and then used it to calculate distances from it for all species in the dataset using the common interval score in CREx (Bernt et al. 2007). This is a similarity measure, where high values indicate highly similar gene orders and vice versa, so negative correlation values with other parameters imply a positive correlation with GORR and vice versa. We used the *AnalyzeCodonData*.*bf* function in HyPhy (Kosakovsky Pond et al. 2020) to infer *ω* values (dN/dS) on a dataset comprising concatenated 13 PCGs of all mitogenomes, with the best-fit GY codon model selected by ModelFinder (Kalyaanamoorthy et al. 2017). Relaxation of selective pressures in selected lineages was further tested using the RELAX (Wertheim et al. 2015) function in HyPhy. The IQ-TREE GHOST algorithm was used to analyse the dataset divided into classes of sites according to their evolutionary background (Crotty et al. 2020). *N*_*e*_ estimation was conducted exactly as described before (Jakovlić et al. 2021), so details are given in Supplementary file S1.

For standard statistical analyses, we used Pearson’s correlation (two measurement variables), Spearman’s correlation (ranked + measurement), and Polyserial correlation (category + measurement). As our data violated the assumption of independence, we used ANOVA phylogenetic generalised least squares (PGLS) in the *nlme* package to conduct pairwise and multigroup comparisons (Pinheiro et al. 2017). We also conducted multilevel regression analyses using two different models that can remove the effect of the evolutionary relationships of species when fitting a regression between variables: *brms* (Bürkner 2018) and *lmekin* (Therneau 2018). For these, we used a matrix of phylogenetic distances extracted from the tree inferred using the CAT-GTR model using the *ape* package. Life history categories (ectotherm, endotherm, and non-parasitic) were divided into pairs of two dichotomous variables and recoded using dummy coding (0 and 1) to test the impact of parasitism (all species divided into parasitic and non-parasitic categories) and thermic habitat (ectothermic vs. endothermic host, with free-living removed from the analyses).

## Results

We used mitogenomes of 223 flatworm species to infer their phylogeny, branch length, maximum longevity, GC skew, GORR, and genome length. These data were used to conduct a series of statistical analyses aimed at testing our working hypotheses. For standard statistical analyses, we categorised all species along the phylogeny+parasitism lines into free-living and parasitic, and the latter (Neodermata) were further subdivided along the taxonomic class lines (‘phylogenetic dataset’). For the ‘thermic dataset’, parasitic species were categorised according to the thermic environment of their intermediate and definitive hosts into ectotherms and endotherms (Fig. 1). For the phylogeny-corrected multivariate analyses, we had to dummy-code the categories, so we tested two datasets: for the ‘parasitism dataset’ all species were divided into parasitic (1) and free-living (0); for the ‘thermic dataset’, free-living were not used, and endotherms and ectotherms were coded as 1 and 0 respectively. This way we also tested for the existence of incongruent signals between lineages, as the latter dataset contains only Neodermata, and the former includes the basal non-parasitic radiation, which exhibits very different evolutionary patterns.

### Hypothesis 1: Functional constraints (thermic habitat and sequence evolution)

We used the ‘thermic’ dataset to test the hypothesis that the thermic environment of the host may influence the mitogenomic evolution of the parasite. Branch lengths were marginally, but statistically significantly, longer in endotherms than in ectotherms (Fig. 2K and L, Supplementary file S2: table S4). GC skews and ω values were also higher in endotherms (significantly and nonsignificantly respectively; Fig. 2I and J). Results were almost identical when the dataset was divided according to the intermediate host (Supplementary file S1). Multivariate analyses indicated that thermic type was the third-best (after GORR and GC skew), but a nonsignificant, predictor of branch length, explaining less than 2% of the branch length variability (Table 2). Remarkably, the thermic host environment and branch length were consistently the best predictors of the GC skew magnitude. RELAX analyses revealed that selection is highly significantly intensified in ectotherms and relaxed in endotherms (both p=0.0000; both compared to all other lineages in the dataset). In conclusion, we found multiple indications in support of the working hypothesis, but the overall relative impact of thermic habitat on mitogenomic evolution seems to be very limited in magnitude.

**Fig. 2.**
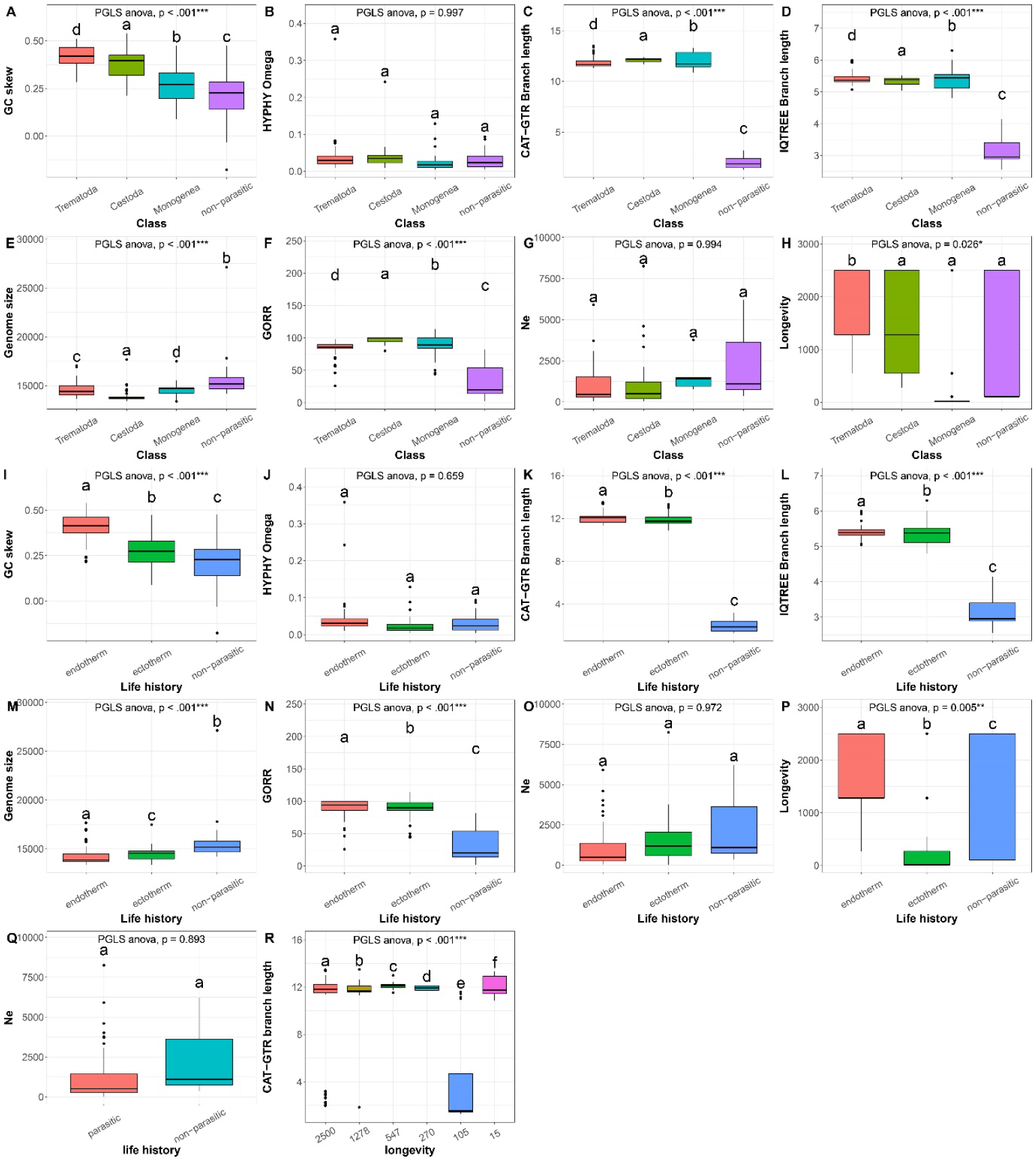
Statistical comparisons of different parameters inferred using mitogenomes of flatworms. Neodermata were grouped in two ways: by taxonomic lines (Class) and with respect to the thermic type of the definitive host (Life history). In the ‘longevity’ dataset, all flatworms were grouped into six categories according to their maximum life span. The parameter compared is shown on the y-axis. GORR is the gene order rearrangement rate (0 indicates no similarity). *N*_*e*_ is the effective population size. Omega is the ratio of nonsynonymous to synonymous mutations (*ω* = dN/dS). PGLS ANOVA parameter shows the result of ANOVA analysis controlled for phylogenetic relationships among the species using *nlme*. *p<0.05, ***p<0.001. Pairwise statistical comparisons between pairs of groups were also conducted using PGLS ANOVA controlled for phylogenetic relationships. The statistical significance (p<0.05) of pairwise comparisons is indicated by different letters above box plots.

**Table 2.**
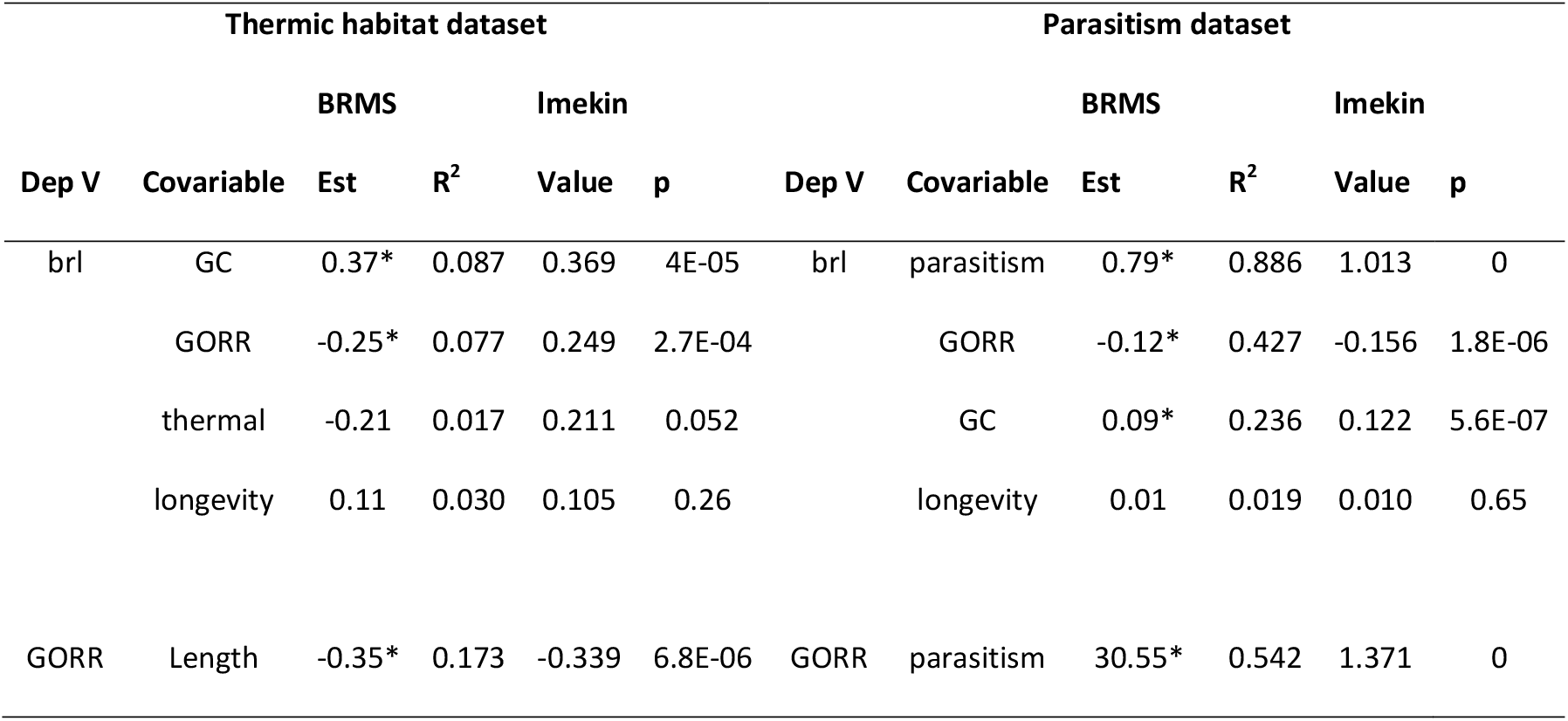
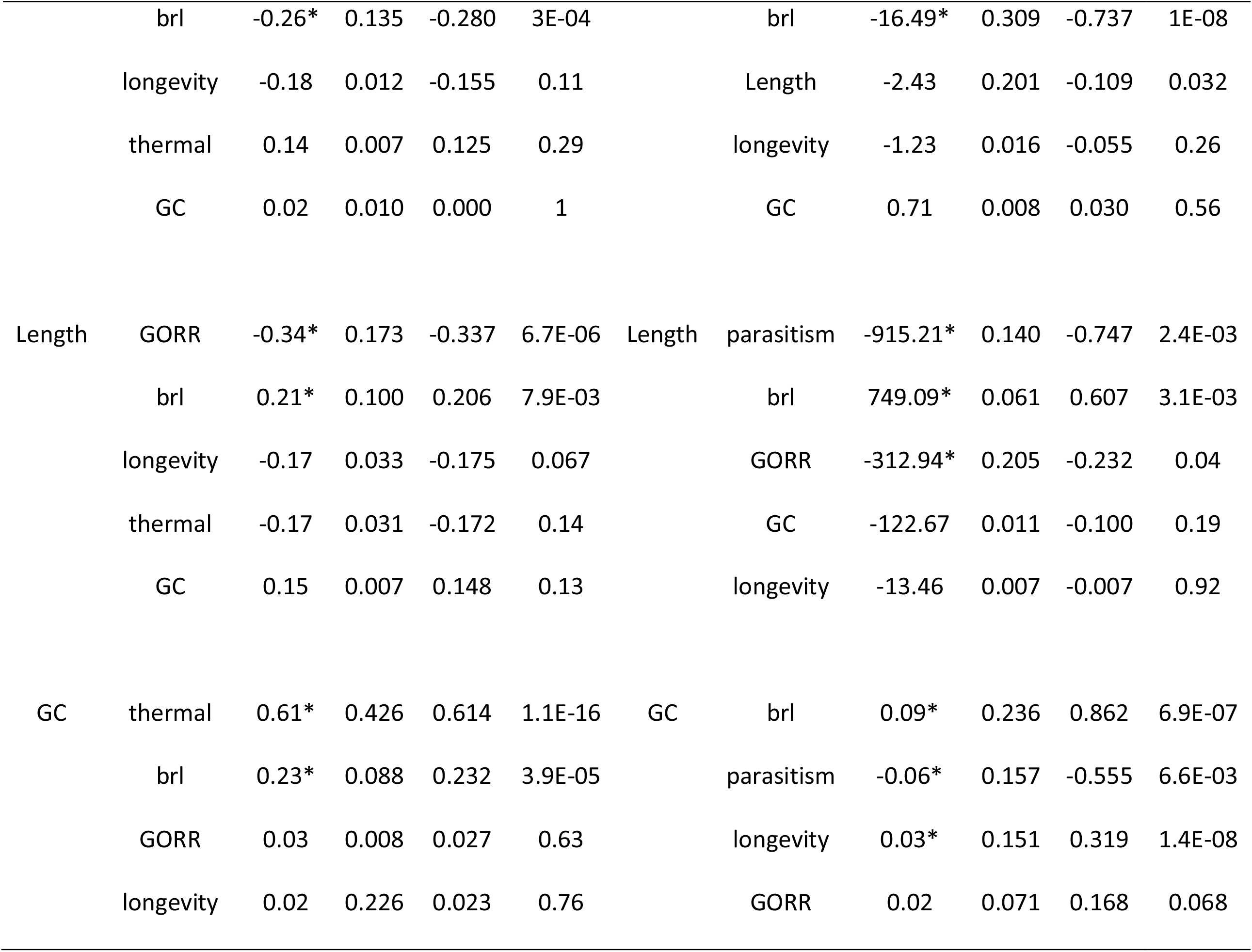
Multilevel regression results inferred using phylogenetic multilevel models implemented in *brms* and *lmekin*. In the parasitism dataset, all species are divided into parasitic and free-living, and in the thermic habitat dataset, all parasitic species are divided into endotherms and ectotherms by the definitive host type. Results for the brl (branch length) and GC (GC skew) variables were calculated using the full dataset, whereas results for the length (mitogenome length) variables were calculated using a dataset containing only putatively complete mitogenomes. Dep V is the dependent variable. Est and Value are estimated effects of covariables on the Dep V, where R2 (BRMS) is the proportion of the Dep V variability explained by the covariable, and p indicates the statistical significance of the impact (*lmekin*). * symbol in the BRMS Est. value column indicates a significant effect of covariable (two-sided 95% Credible intervals do not span 0). Full data are available in Supplementary file S2: tables S1 and S2. GORR: gene order rearrangement rate.

### Hypothesis 2: Race for replication (thermic habitat and genome size)

In weak support of the ‘race for replication’ hypothesis, genome size was marginally but significantly smaller in endotherms (14.2 Kbp) than in ectotherms (14.5 Kbp). It was by far the largest in non- parasitic species (16.1 Kbp) (Fig. 2M). Multivariate analysis with genome length as the dependent variable indicated that the thermic environment had a minor, nonsignificant, impact, explaining ≈3% of the mitogenome size variability. Contrary to the predictions of the ‘race for replication’ hypothesis, the ectotherm size range (13.4 to 17.5 Kbp) was slightly narrower than the endotherm range (13.4 to 17.7 Kbp). Non-parasitic exhibited by far the widest range: 14.2 to 27.1 Kbp. Although we found weak support for some predictions of the ‘race for replication’ hypothesis, overall data indicate that its effects on mitogenomic evolution are at best minimal.

### Hypothesis 3: gene order and mitogenome size

Mitogenomes were significantly smaller in cestodes (13.8 Kbp) than in trematodes and monogeneans (≈14.6 Kbp), and the largest in non-parasitic lineages (16 Kbp), which perfectly matched the GORR pattern (Fig. 2E and F). Indeed, the genome size was significantly positively correlated with GORR (−0.45, negative values imply positive correlation; Fig. 3). In multivariate analysis of the thermic dataset with GORR as the dependent variable, mitogenome size was the best predictor with the largest effect; and in the analysis of the parasitic dataset, it had a significant but small effect (parasitism and branch length had much greater effects; Table 2). With mitogenome length as the dependent variable, results were similar, but GORR did not have such consistently significant effects. The association was positive (negative values) in both cases, and the explanatory power (mitogenome size on gene order variability and vice versa) was consistent in all four analyses: roughly around 20% (Table 2).We can conclude that this dataset supports hypothesis 3.

**Fig. 3.**
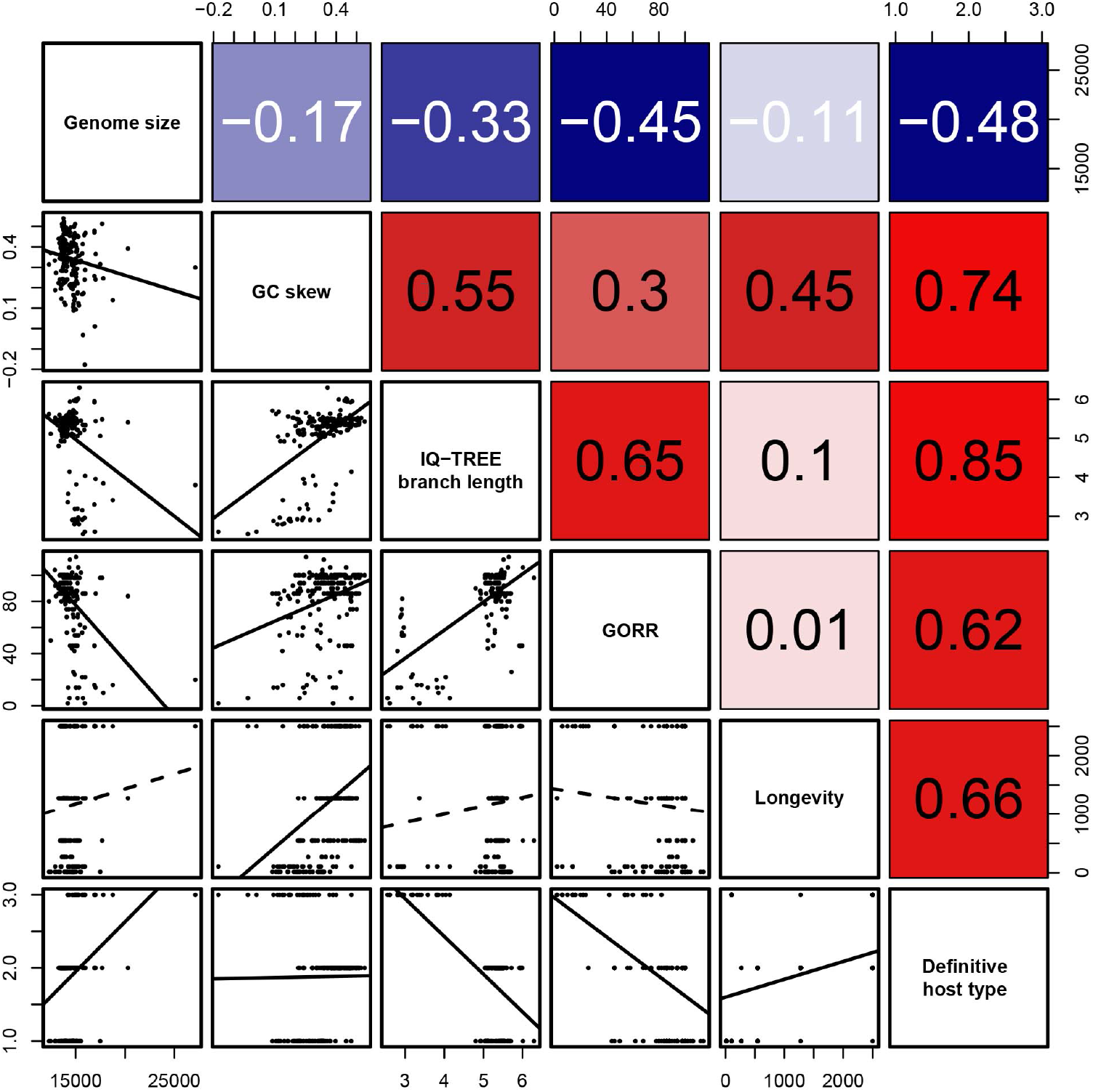
Correlation analyses. Above-right of the diagonal are correlation values, also shown as a correlation heatmap (0 = grey; 1 = red, -1 = blue). Below-left of the diagonal are data plots, where dotted/solid trendlines represent nonsignificant/significant (P>0.05/P<0.05) correlations respectively.

### Hypothesis 4: Gene order and sequence evolution

Gene order rearrangements were negatively correlated with branch length (0.65; Fig. 3). Non- parasitic species (shortest branches) had by far the most rearranged gene orders, whereas the remaining three classes had comparatively similar average values (Fig. 2F; low values = high rearrangement rate, which also means that positive values imply negative correlation in statistical analyses). In multivariate analysis with branch length as the dependent variable, GORR was the 2^nd^ best predictor, but the R^2^ parameter indicates that the effect was much greater in the parasitism dataset than in the thermic dataset (8% vs. 43% respectively; Table 2). With GORR as the dependent variable, branch length was a strong and highly significant predictor in both datasets, explaining 14% and 31% of the variability (Table 2). However, contrary to the negative correlation in the standard statistical analysis, the Est. values were negative (implying a positive correlation) in all multilevel regression analyses. We tested whether this may have been an artefact caused by the collinearity of independent variables: with GORR as the only independent variable, its Est. value was weakly positive (0.02). As this only partially explains the discrepancy, we further tested whether there is strong interlineage variability in these signals. Indeed, the correlation between branch length and GORR within the Neodermata was negative (−0.26, p = 0.0002; Supplementary figure S2). This explains the discrepancy between the results: non-parasitic species possess by far the most rearranged mitogenomes and shortest branches (negative correlation), but within the Neodermata the correlation between the two variables is weakly but significantly positive.

### Hypothesis 5: Longevity-dependent selection

Longevity did not exhibit any correlation with the branch length (0.1; Fig. 3). This absence of correlation was reflected in the absence of a clear trend in branch length among different longevity categories, with by far the shortest average branch lengths exhibited by the relatively short-living category 2 (comprising most of the free-living species; Fig. 2R). Along the taxonomic lines, longevity was the highest in Trematoda, followed by Cestoda, non-parasitic, and finally Monogenea (Fig. 2H). Along the thermic habitat lines, endotherms exhibited the highest longevity, followed by non-parasitic and finally ectotherms (all differences were significant; Fig. 2P). In multivariate analyses, longevity had a very weak, nonsignificant, effect on the branch length, explaining only 2-3% of the variability, in both datasets (Table 2).

### Hypothesis 6: Parasitism

To test this hypothesis, we used the ‘phylogenetic dataset’. Although branch lengths were much greater in the CAT-GTR analysis than in the IQ-TREE analysis, the two analyses produced relatively congruent results in terms of relationships among different lineages, so branch lengths were highly correlated (p<0.05; 0.97) between the two methods (Supplementary figure S1). The three neodermatan groups (Monogenea, Cestoda and Trematoda) had very similar average values (although differences were highly significant): 11.90 – 12.09 in CAT-GTR, and 5.33 – 5.43 in IQ-tree (Fig. 2C and D). Branches were much shorter in non-parasitic species in both analyses (1.96 in CAT-GTR, and 3.17 in IQ-TREE). GC skews were the lowest in non-parasitic lineages (0.21), and they exhibited a significant correlation with the branch length (0.55; Figs. 2 and 3). For the phylogeny- corrected multivariate analyses, parasitism was the best predictor of branch length, explaining 89% of the variability (several other variables also had significant effects; Table 2; Supplementary file S2: table S1). These findings are in agreement with the working hypothesis and indicate either that parasitic lineages evolve under relaxed purifying selection pressure or under elevated mutation pressure. In support of the first scenario, the RELAX analysis revealed that selection is highly significantly intensified in non-parasitic lineages, and relaxed in parasitic lineages (both p<0.0001; both compared to each other). To further attempt to disentangle purifying selection pressures from mutational pressures, we divided the aligned protein-coding genes sequences dataset into 7 classes of sites according to the rate at which they evolve, approximately in the order from slowest – class1 (comprised almost completely of slow-evolving 2^nd^ and 1^st^ codon sites) to fastest - class 7 (comprised almost completely of fast-evolving 3^rd^ codon sites) using the GHOST algorithm. Classes 2 to 5 exhibited similar patterns to the overall dataset: by far the shortest branches in non-parasitic and similar branch lengths in the other three groups (Supplementary file S1). Classes 1 and 6 also produced the shortest branches for non-parasitic, but Monogenea, Trematoda and Cestoda differed from each other. Class 7 (90% 3^rd^ codon sites) exhibited an almost fully inverted pattern: non-parasitic > Trematoda > Monogenea > Cestoda. However, the non-parasitic lineages had a very wide distribution of data. As class 7 sites presumably evolve under the weakest purifying selection pressure, this finding indicates that non-parasitic species may be evolving under stronger mutational pressures due to their higher metabolic rates, but overall they evolve slower than other lineages because of stronger purifying selection pressures.

### Hypothesis 7: The effective population size (*N*_*e*_)

We attempted to assess whether the elevated evolutionary rates in parasitic flatworms are attributable to reduced *N*_*e*_, but the resolution of our analysis was weakened by the fact that we only managed to obtain *N*_*e*_ data for 64 endotherms, 8 ectotherms, and 5 non-parasitic species. The faster-evolving endotherm lineages had much lower average *N*_*e*_ values than ectotherm lineages (1044 vs. 2093), but the values did not differ significantly due to wide *N*_*e*_ value distribution in endotherms (Fig. 2O). Similarly, the faster-evolving parasitic lineages had much lower average *N*_*e*_ values than free-living lineages (1160 vs. 2412), but the values did not differ significantly due to the wide *N*_*e*_ value distribution in parasites (Fig. 2Q). In multivariate analyses, *N*_*e*_ values did not have a significant effect on branch lengths, or any other variable (Supplementary file S2). Overall, we did not find support for this hypothesis, but a stronger dataset is needed to test it thoroughly.

### Hypothesis 7: Relative impacts of different variables

We found evidence that multiple factors simultaneously impact mitogenomic evolution in flatworms, but their relative contributions were very different. For example, branch lengths were greater in endotherms than in ectotherms, but the magnitude of difference (5.4 vs. 5.35 in the IQ-TREE) was almost negligible compared to the difference between parasitic and non-parasitic lineages (5.38 vs. 3.17, Fig. 2K and L, Supplementary file S2: table S4). This was confirmed by multivariate analyses, where both variables had a significant impact on the branch length, but whereas the impact of parasitism explained almost 90% of the branch length variability, thermic habitat impact explained less than 2% of the variability (Fig. 2, Table 2).

## Discussion

We tested predictions of eight different hypotheses in relation to mitogenomic evolution (both sequence and architecture) in flatworms. It should be noted that some of these hypotheses have contradictory predictions. For example, ‘functional constraints’ and ‘race for replication’ hypotheses predict that the impact of thermic habitat and metabolic rate on the mitochondrial evolution should have almost opposite effects on the primary sequence evolution (faster evolution in endotherms) and genome size and complexity (smaller and less size-variable mitogenomes in endotherms) (Rand 1994). Moreover, whereas hypotheses 1 and 2 together predict that mitogenomic sequence evolution and mitogenomic complexity (size) will be negatively correlated in endotherms (higher and lower respectively), hypotheses 3 and 4 predict that GORR, mitogenome size, and branch length should all be positively correlated. This makes it impossible to find support for all tested hypotheses.

### Sequence evolution is faster in parasites associated with the thermally stable environment of endothermic hosts, but the effect is small

The influence of thermic habitat has received some attention in the early mitogenomic evolutionary studies (Thomas and Beckenbach 1989; Rand 1994), but it has been only sporadically revisited by later studies (Lagisz et al. 2013; Lajbner et al. 2018). We found multiple indications in support of this hypothesis: longer branches, higher GC skews, and higher *ω* values in endotherms. However, thermic habitat was a very weak predictor of branch length, as evidenced by multivariate analyses and only marginally different branch lengths. Our results closely correspond to those in nematodes: (Lagisz et al. 2013) found some support for the functional constraints hypothesis, but the evidence was not sufficiently strong to accept it with confidence. We further assessed whether intermediate host type affects the results, but we obtained similar results. In conclusion, there is some evidence that thermally stable environments of endothermic hosts may allow for slightly relaxed purifying selection pressures in their flatworm parasites in comparison to parasites of ectothermic hosts, but the effect is relatively minor.

### Mitogenomes are smaller in parasites associated with the thermally stable environment of endothermic hosts, but the size range is wider

In partial agreement with the ‘race for replication’ hypothesis, which proposes that mitogenomes should be smaller in size and exhibit reduced size variability in endotherms (Rand 1993), including their parasites (Lagisz et al. 2013), mitogenomes were smaller in flatworm parasites associated with the thermally stable environment of endothermic hosts. Contrary to these predictions, the endotherm range was slightly wider than the ectotherm range. Therefore, the results are too weak and inconsistent to accept the hypothesis with confidence (additional discussion in Supplementary file S1).

### Mitogenomic architecture rearrangements are (usually) positively correlated with mitogenomic size

Along with the fact that some types of rearrangement events produce an increased abundance of noncoding segments between genes (Boore 2000), we hypothesised that mutations in the replication and maintenance machinery should result both in increased gene reshuffling and sequence duplication errors, from which we inferred that GORR should be positively associated with the mitogenome size. Surprisingly, aside from the positive correlation between the mitogenome length and GORR established in nematodes (compact mitogenomes were more structurally stable) (Lagisz et al. 2013), the correlation between mitogenomic size and other relevant parameters does not appear to have been tested statistically in other studies to our knowledge. Previous studies have observed that exceptionally large mitogenomes in nematodes are often associated with large noncoding regions, segmental duplications, and hypervariable structure (Hyman et al. 2011; Zou et al. 2017), but a recent study found that nematode lineage Longidoridae contradicts the prediction that compact mitogenomes should be structurally stable (Zou et al. 2022*a*). In conclusion, although we found strong evidence for our hypothesis, there is evidence that the association between the two variables is not universal, and we should also account for the possibility that the reliability of mitogenome size analyses may be affected by sequencing artefacts.

### The correlation between mitogenomic architecture rearrangements and sequence evolution is inconsistent among lineages

The causes of mitochondrial gene rearrangements remain unclear. Previous studies suggested that there may exist a correlation between gene rearrangements and a variety of external forces, including elevated nucleotide substitution rates (Shao et al. 2003; Xu et al. 2006; Bernt et al. 2013; Zou et al. 2022*b*), the adoption of parasitic lifestyles (Dowton and Austin 1995; Oliveira et al. 2008), or the inhabitation of ‘extreme’ ecological niches (e.g. deep-sea, cold waters, hydrothermal vents) (Zhuang and Cheng 2010; Gan et al. 2018). However, we suspect that there is no causative relationship behind any of these correlations. It was proposed that genomic mutations causing a decrease in the fidelity of mitogenome replication are a major cause of high rates of genome rearrangement as well as point mutations (Xu et al. 2006) and that gene order rearrangements should be affected by the purifying selection as they may affect the regulation of gene expression (Boore 1999). Reduced purifying selection pressures and their association with *N*_*e*_ would also be a suitable explanation for the above-mentioned associations (mutation rate, parasitism, extreme niches, etc.), which implies that architectural and sequence evolution rates should be correlated.

Contrary to this hypothesis and the above evidence, our findings indicate that gene order rearrangements and sequence evolution are almost completely decoupled in non-parasitic flatworms. Specifically, in the Neodermata, the correlation is weakly but significantly positive, but non-parasitic lineages exhibit highly rearranged mitogenomes and short branches. Intriguingly, Bernt et al. observed that in Bilateria substitution rate is positively correlated with GORR, but argued that this association does not fully explain the exceptionally high substitution rates in Platyhelminthes (Bernt et al. 2013). Similarly, there was no correlation between gene rearrangement events and species ecology or lineage-specific nucleotide substitution rates in Decapoda (Crustacea) (Tan et al. 2019). These results indicate that the relationship between GORR and sequence evolution is random and lineage-specific within the Platyhelminthes. Overall, we can conclude that for currently unknown reasons purifying selection pressure does not ffect architectural rearrangements equally in all lineages. More studies, encompassing a broad range of metazoan lineages, are needed to better understand the relationship between these two variables.

### Longevity does not have a consistent impact on mitogenomic evolution in flatworms

Parasite longevity is expected to be inversely correlated to the thermic environment of the host: at higher temperatures parasites invest in higher metabolic and reproductive rates, but a shorter life span appears to be an unavoidable tradeoff (Bakke et al. 2007). Our analyses indicate that longevity has no significant effect on the branch length in flatworms, but it does have a significant effect on GC skew and GORR. It remains unknown whether these correlations are spurious, or whether they may exist an underlying relationship between these variables. Our study provides further evidence that the impact of longevity may be strongly lineage-specific in metazoans (Thomas et al. 2010; Allio et al. 2017) (see further discussion in Supplementary file S1).

### Parasitic flatworm lineages exhibit higher evolutionary rates than non-parasitic lineages

We found strong support for the hypothesis that parasitic lineages evolve faster in flatworms. The difference was striking, as the branches of non-parasitic species were almost six times shorter than those parasitic lineages in the CAT-GTR analysis, and 1.5 times shorter in the IQ-tree analysis (Fig. 1). Multivariate analyses also support this, as life history categorisation into parasitic and free-living lineages (called ‘parasitism’ in the analyses) was by far the best predictor of branch length (Table 2). Previous evidence that parasitism might be associated with elevated mitochondrial sequence evolution rates is mostly confined to some arthropod lineages (Dowton and Austin 1995; Castro et al. 2002; Shao et al. 2003; Hassanin 2006; Shao and Barker 2007; Oliveira et al. 2008; Bernt et al. 2013; Jakovlić et al. 2021), so our study provides important evidence that this association may not be lineage-specific.

It has been proposed that possible causes for this correlation between parasitism and elevated evolutionary rates might be increased flux of mutagens (Dowton and Campbell 2001), the compensation-draft feedback (Oliveira et al. 2008), or *N*_*e*_ reductions caused by high speciation rates and frequent founder events during transmissions to new host individuals (Page et al. 1998; Castro et al. 2002). We did find evidence for relaxed purifying selection in parasitic lineages and a statistically nonsignificant reduction in the population size in parasitic lineages, but *N*_*e*_ did not feature as a key factor influencing the mitogenomic evolution in any of our multilevel regression analyses. Several studies found that *N*_*e*_ is not a reliable predictor of mitogenomic evolution (Bazin et al. 2006; Jakovlić et al. 2021), but our *N*_*e*_ analyses were weakened by the limited availability of molecular data for many lineages, which may have affected both the precision of our *N*_*e*_ estimates and subsequent multilevel regression analyses (further discussion in Supplementary file S1). Our analyses were further hampered by the fact that parasitic flatworm lineages included in our study are monophyletic. This means that we cannot exclude a possibility of a deleterious mutation in the mitogenomic replication and quality control mechanism occurring in the common ancestor of Neodermata. However, it would be much easier to explain such a scenario in the context of pre-existing relaxed purifying selection pressures in parasitic lineages, as well as the phylogenetic correction algorithm we used, so it does not preclude the key role of parasitism.

A recent study proposed that this association between parasitism and elevated evolutionary rates could be driven by the reduction of locomotory capacity in parasitic lineages (Jakovlić et al. 2021). Because free-living flatworms have to search for food and evade predators, they are expected to have much stronger locomotory capacity than parasitic flatworms, which rely on the host to provide food and evade predators. However, compared to many highly locomotory free-living crustacean lineages, the locomotory capacity of free-living flatworms is very weak, so flatworms exhibit much smaller interlineage variability than crustaceans in this variable. As a result, we must account for the possibility that the reduction of purifying selection pressures in parasitic species may be a result of multiple other factors, including the locomotory capacity, *N*_*e*_ magnitude, and the overall reduction of various biological processes in parasites (Keeling et al. 2010), which might allow additional reduction of energy production and metabolic rates. Remarkably, GHOST analyses indicated that non-parasitic species may be evolving both under the strongest purifying selection (GHOST class 1) and mutational pressures (GHOST class 7), possibly due to their higher metabolic rates. This would support a model in agreement with observations in sea animals, where locomotory capacity is positively correlated with the metabolic rate (Seibel and Drazen 2007). Therefore, multiple factors may act synergistically to produce this relationship, but their relative impacts appear to be somewhat stochastic and thus inconsistent and lineage-specific. In conclusion, our analyses support previous observations that evolutionary rates are higher in mitogenomes of parasites, but due to the monophyletic origin of parasitic lineages in flatworms, we cannot exclude the impact of other variables.

### Multifactorial mitogenomic evolution: inconsistency, interdependence, and magnitude of effects

Mitogenomic evolutionary studies are plagued by inconsistent and contradictory findings and lineage-specific effects, which suggests that there is probably no universal explanation for the mitogenomic evolution patterns in metazoans (Bazin et al. 2006; Allio et al. 2017; Jakovlić et al. 2021). Most early findings were inferred using vertebrate datasets, but when later studies started incorporating invertebrates, it became clear that what seemed to be consistent correlations might actually be vertebrate-specific effects. For example, whereas the associations between mitogenomic evolution and body size and longevity are relatively well-established in vertebrates, the evidence in invertebrates is inconsistent (Nabholz et al. 2008, 2016; Thomas et al. 2010; Saclier et al. 2018; Jakovlić et al. 2021). Indeed, our results show that correlation values between parameters vary notably depending on whether the basal radiation of non-parasitic flatworm lineages is included or not (Supplementary figures S1 and S2), but conclusions were affected only for the correlation between GORR and branch lengths variables. We argue that a major factor causing the inconsistency of results, which seems to have been overlooked by many previous studies, is the varying magnitude of effects of different variables among lineages. Many previous studies focused on determining whether a certain factor influences the mitogenomic evolution in a certain lineage, but this may produce inconsistent results when multiple factors are not simultaneously accounted for.

Furthermore, the magnitude of the impact of a specific variable may be directly positively associated with the degree of intralineage variability in that variable. For example, almost all bird species are evolving under relatively strong selection for locomotory capacity, so this variable may have smaller effects than more variable factors, such as body size or *N*_*e*_ (Lanfear et al. 2010; Thomas et al. 2010; Nabholz et al. 2016); contrary to this, in lineages that exhibit extreme variability in locomotory capacity, such as crustaceans, this variable may almost completely obscure the signal from variables such as body size *N*_*e*_, thus causing inconsistent findings among studies (Jakovlić et al. 2021). In flatworms, both thermic habitat and (putatively) parasitism have significant impacts on the evolutionary rate, but strong effects of the second parameter obscure the signal produced by the former parameter.

### Conclusions

Our analyses show that multiple factors affect the mitogenomic evolution in flatworms, but there is a strong variability in the magnitude of their effects. The greatest effect appears to be generated by the parasitic lifestyle, which expands the previous indications that parasitism might be associated with elevated mitochondrial sequence evolution rates in flatworms, but our analyses were hampered by the monophyletic origin of parasitic flatworm lineages, so we cannot attribute the observed effect to parasitism with confidence. In general, the non-parasitic lineages exhibit several unique features within the flatworm dataset, which makes them very interesting from the evolutionary perspective: by far the slowest sequence evolution and by far the highest gene order rearrangement rate. Contrary to many previous findings, this indicates that these two variables are not consistently correlated in metazoans. The picture that emerges is that gene order rearrangements are strongly selected against in some lineages, whereas in some lineages most rearrangements seem to be weakly affected by the purifying selection pressures (Xu et al. 2006; Zou et al. 2017), which results in a spurious nature of the correlation between the gene order rearrangements and sequence evolution. In conclusion, our results partially confirm previous findings that patterns of mitogenomic evolution are almost unpredictable across different metazoan lineages (Bazin et al. 2006; Allio et al. 2017), and we argue that the underlying reason for this unpredictability is the multifactorial nature of mitogenomic evolution, with lineage-specific dominant factors. More specifically, the strong interlineage variability in the magnitude of effects of multiple factors that affect the mitogenomic evolution produces strong lineage-specific mitogenomic evolution patterns.

## Acknowledgements

We would like to thank Dr. Xiang Liu for helpful discussions and technical assistance.

## Funding sources

This work was supported by the National Natural Science Foundation of China (32102840, 31872604); the Start-up Funds of Introduced Talent in Lanzhou University (561120206). The funders had no role in study design, data collection and analysis, decision to publish, or preparation of the manuscript.

### Data access

All data used in this study were retrieved from the NCBI’s GenBank database. A detailed overview of data, with GenBank accession numbers, can be found in Supplementary file S2: table S3.

