## Supplementary file S1 for "Drivers of interlineage variability in mitogenomic evolutionary rates in flatworms (Platyhelminthes) are multifactorial"

**Drivers of interlineage variability in mitogenomic evolutionary rates in flatworms (Platyhelminthes) are multifactorial**

### Supplementary Methods

#### Datasets

A subset of 202 species was used for multivariate analyses after removing all species with dN = 0, no GORR, and no host type. For pairwise comparison analysis, we used several subset datasets, depending on the focus of the analysis: (1) removing species with $\omega = 0$ or 999 (indicate infinity) resulted in 217 species; (2) removing species for which we failed to infer the gene order distance (too many missing genes) or the distance was 0 resulted in 220 species; (3) removing species which contain less than 36 genes (to evaluate the effect of missing genes) resulted in 171 species; (4) removing species exhibiting incomplete mitogenomes resulted in 182 species; (5) removing species for which we failed to infer the $N_e$ and those with $N_e = 0$ resulted in 77 species. As we could not be certain that we removed all incomplete mitogenomes, we applied two different thresholds for the removal of mitogenomes and conducted analyses with both. Specific numbers varied, but conclusions were unaffected, so we opted to use the larger dataset, to maximise the amount of data. For GC skews, noncoding segments were excluded to minimise the noise caused by varying NCR ratios. For a few non-parasitic species that had genes encoded on both strands (flatworms general have all genes encoded on a single strand), we used the complementary strand of the coding section to calculate GC skews (so if a gene was encoded on the minus strand, we used the complementary plus-strand sequence to calculate skews).

#### Phylogenetic analyses

As the PhyloBayes analysis for which we used the nucleotide (NUC) dataset did not fully converge, we also run the same analysis using the amino acids (AAs) dataset. This analysis fully converged (maxdiff < 0.1 and minimum effective size > 300). The topologies of IQ-TREE and PhyloBayes were highly congruent, and branch lengths exhibited an almost perfect correlation of 0.98). On this basis, we used the NUC tree for subsequent analyses.

#### Multivariate regression

In the *brms* analysis, every parameter was summarized using the mean (Est.) and Bayesian $R^2$ (Est.) values, proportional to the importance of individual predictors. Bayesian $R^2$ (Est.) is calculated via univariate regression and evaluates how much a variable can explain the variance of the dependent variable. In most cases, the mean estimate and Bayesian $R^2$ were in agreement. In such cases interpretation was straightforward. In cases where the signal varied between the two parameters, we also accounted for confidence intervals (CI lower and upper 95%). When CI spanned 0 (i.e.

ranged from negative to positive values), we regarded this result as unreliable and chose the CI range that did not span 0 as the better explanatory variable.

### dN/dS analyses

The rate of nonsynonymous (dN) to synonymous (dS) substitutions ($\omega$ = dN/dS) is a commonly used indicator for measuring the direction and magnitude of selective pressure on genes, where $\omega$ is inversely correlated to the strength of purifying selection. As it is difficult to calculate these parameters with confidence, we used two different methods and a dataset comprising concatenated 13 PCGs of all mitogenomes: AnalyzeCodonData.bf function in HyPhy (Kosakovsky Pond et al. 2020) with the best-fit GY codon model selected by ModelFinder (Kalyaanamoorthy et al. 2017). It should be noted that the inference of $\omega$ patterns is reliable only for testing specific hypotheses. Due to them being very parameter-rich, they are prone to producing unreliable results when inferring absolute dN/dS ratios. The divergence of neutral sites is proportional to mutation rate, so *dS* can be used as an imperfect proxy for estimating the strength of the mutation rate; *dN* is expected to be influenced both by the mutation rate and by population size; and $\omega$ is expected to be influenced by selection and effective population size, but not the mutation rate (Kimura 1968; Lanfear et al. 2010).

### $N_e$ estimation

We mined the BOLD database (Ratnasingham and Hebert 2007) for *cox1* data for the 223 flatworm species (corresponding to the mitogenomic dataset) using the *bold* package in R (Chamberlain 2020). After removing species with less than two *cox1* entries, the dataset contained 89 species. The formula $\Theta_W = 2N_e\mu_{bg}$ was used to infer $N_e$, where $\mu_{bg}$ is the mutation (or substitution) rate per site per million years, and $\Theta_W$ is Watterson's theta estimator for a scaled mutation rate (Watterson 1975). Sequences were aligned in batches using the MAFFT plugin in PhyloSuite and used to calculate lineage-specific $\Theta_W$ with DendroPy (Sukumaran and Holder 2010). Lineage-specific substitution rates ($\mu_{bg}$) were inferred using the BEAST 1 package (Suchard et al. 2018), with an uncorrelated, relaxed clock model with lognormal distribution and Yule model as the tree prior, and $6\times10^8$ of Bayesian MCMC generations, with sampling every 2000 generations. For calibration points see the main document. Tracer v.1.7 was used to determine when the convergence was reached (effective sampling size > 200) (Rambaut et al. 2018). TreeAnnotator program in BEAST was used to compute the Maximum Clade Credibility (MCC) tree (10% burn-in). Lineage-specific substitution rates were extracted from the resultant tree with the help of the Treeio package (Wang et al. 2020). Finally, lineage-specific $N_e$ values were inferred from the obtained lineage-specific $\Theta_W$ and substitution rates.

### Supplementary Results

#### Hypothesis 1: Functional constraints (thermic habitat and sequence evolution)

Following the hypothesis that the thermic environment of the host may influence the mitogenomic evolution, we analysed the same variables on the 'thermal dataset'. Omega values were the lowest in ectotherms (0.02), followed by non-parasitic (0.03), and the highest in endotherms (0.04). dN and dS values also produced results that did not correspond fully to other variables, as both values were the highest in non-parasitic species, followed by ectotherms, and finally the lowest in endotherms (Figure S1).

#### Hypothesis 6: Parasitism

The HYPHY algorithm resolved Monogenea as evolving under the strongest purifying selection (0.02), followed by non-parasitic (0.03), and Cestoda and Trematoda (both 0.04), but *ω* values did not differ significantly among the four groups (Fig. 2B).

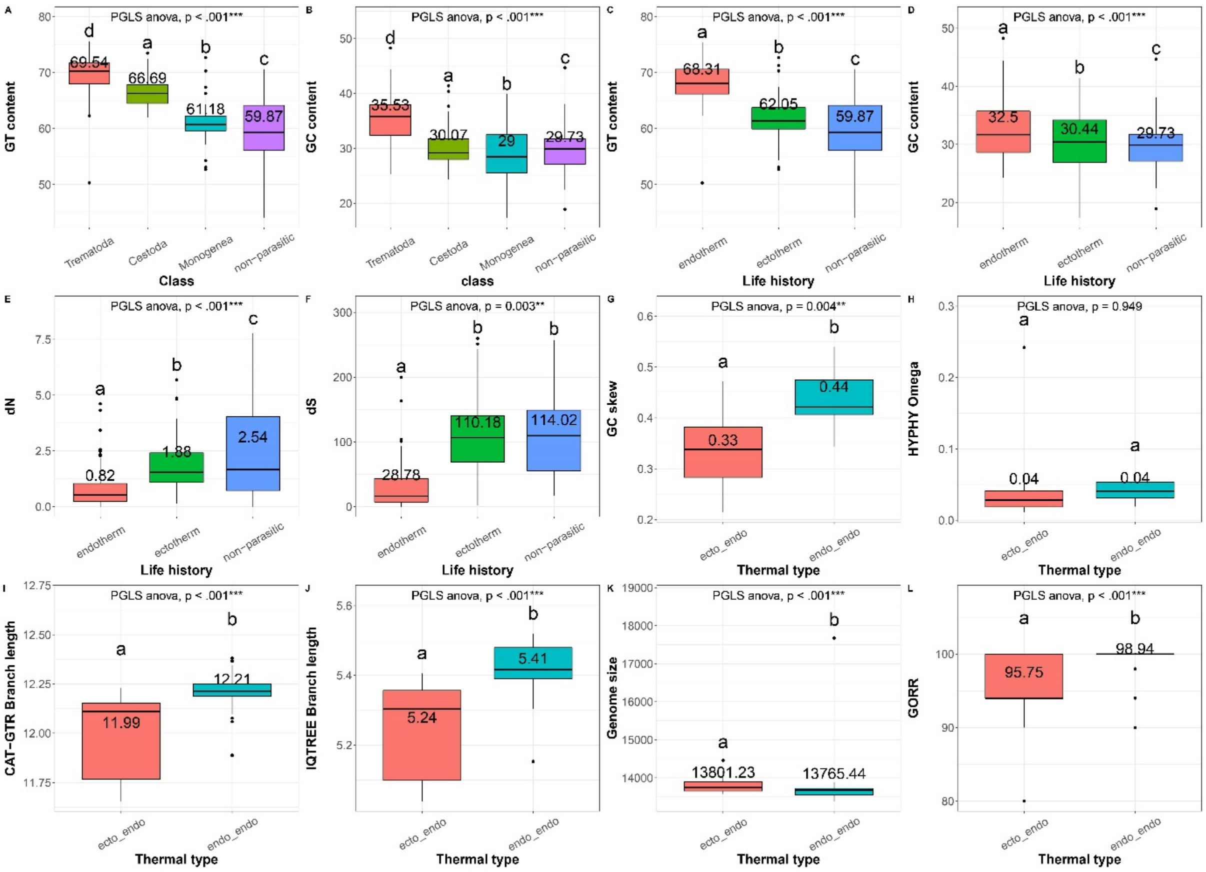

***Figure S1.*** **Statistical comparisons of different parameters inferred using mitogenomes of flatworms.** Neodermata were grouped in two ways: by taxonomic lines (Class) and with respect to the thermic type of the definitive host (Life history). The parameter compared is shown on the y-axis. dS and dN are the numbers of substitutions per synonymous and non-synonymous site, respectively. PGLS ANOVA parameter shows the result of ANOVA analysis controlled for phylogenetic relationships among the species using *nlme*. *$p<0.05$, ***$p<0.001$. Pairwise statistical comparisons between pairs of groups were also conducted using PGLS ANOVA (controlled for phylogenetic relationships). The statistical significance ($p<0.05$) of pairwise comparisons is indicated by different letters above box plots. Text in boxes represents the mean value of each category. Thermal type: thermal type of intermediate and definitive hosts, where ecto_endo indicates ectothermic intermediate host and endothermic definitive host, whereas endo_endo indicates endothermic intermediate and definitive host.

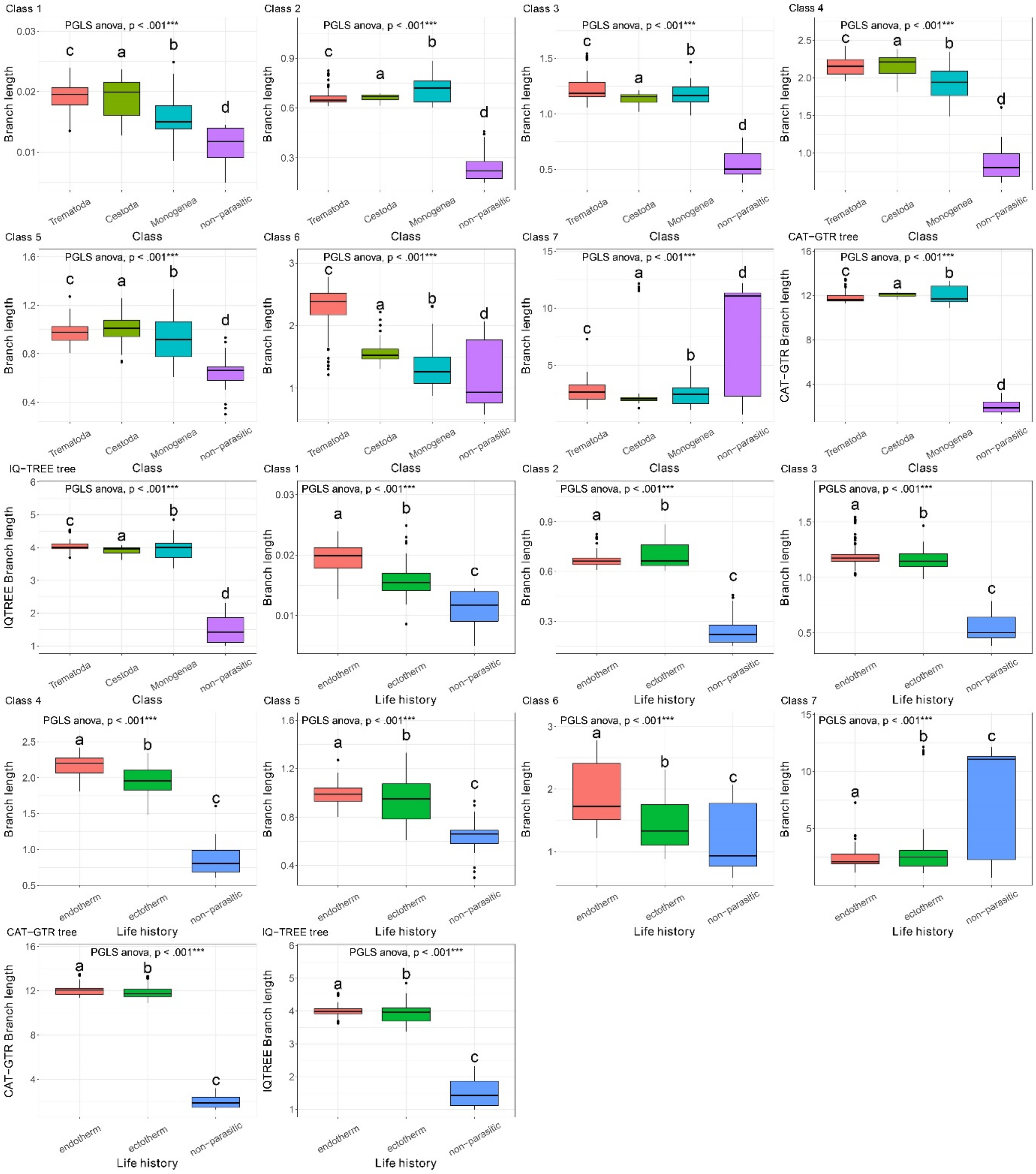

***Figure S2.* Statistical comparisons of GHOST class 1-7.** Neodermata were grouped in two ways: by taxonomic lines (Class) and with respect to the lifestyle and thermic type of the host (Life history). The parameter compared is shown on the y-axis. PGLS ANOVA parameter shows the result of ANOVA analysis controlled for phylogenetic relationships among the species using *nlme*. *p<0.05, ***p<0.001. Pairwise statistical comparisons between pairs of groups were also conducted using PGLS ANOVA. The statistical significance of pairwise comparisons is indicated by different letters above the box plots.

***Table S1.*** The aligned protein-coding genes sequences dataset was divided into 7 classes of sites according to the rate at which they evolve (from slowest – class1, to fastest - class 7) using the IQ-TREE GHOST algorithm. Sites were also classified by the codon position. The numbers represent the proportions of sites.

| Codon | class 1 | class 2 | class 3 | class 4 | class 5 | class 6 | class 7 |
|---|---|---|---|---|---|---|---|
| 1 | 0.3778 | 0.3515 | 0.4217 | 0.6352 | 0.2172 | 0.2410 | 0.0887 |
| 2 | 0.5998 | 0.6308 | 0.5506 | 0.2920 | 0.0610 | 0.0575 | 0.0077 |
| 3 | 0.0222 | 0.0175 | 0.0276 | 0.0727 | 0.7217 | 0.7014 | 0.9035 |

### Supplementary Discussion

#### Sequence evolution is faster in parasites associated with the thermally stable environment of endothermic hosts, but the effect is small

We found that GC content was significantly higher in endotherms (Figure S1), which apparently supports the hypothesis that GC is positively correlated with the environmental temperature (Bernardi 1995). However, several studies (partially) rejected this hypothesis (Hurst and Merchant 2001; Uliano et al. 2010). In our study GC content mirrored the branch length and GC skew patterns (endotherms > ectotherms > free-living), which suggests that mitogenomes of flatworms may simply be evolving towards a higher GC content. Also, the thermic environment was consistently the best predictor of the GC skew magnitude. However, GC content did not mirror branch length and GC skew in the dataset divided along the taxonomic lines, which leaves the issue unresolved.

#### Mitogenomes are smaller in parasites associated with the thermally stable environment of endothermic hosts, but the size range is wider

This finding is logically somewhat perplexing because parasites inhabiting endothermic hosts do not have to produce the thermal energy themselves. The inner workings of the hypothesis propose that higher metabolic demands associated with endothermy impose stronger purifying selection constraints for small genome size (Rand 1993). Indeed, mitogenomes were the largest in non-parasitic species, which we expect to have higher metabolic demands compared to parasites due to their higher locomotory capacity and metabolic requirements. A possible explanation would be that parasites of endothermic hosts invest disproportionately high amounts of energy in reproduction, which results in them having much higher metabolic rates than ectothermic host parasites and non-parasitic flatworms. Unfortunately, comparative metabolic rates of parasites remain poorly understood so this remains conjectural, but there is evidence that parasites have higher fecundity and reduced life span at higher temperatures (Bakke et al. 2007). We should mention here that the putative association between high metabolic rates and mitogenomic size can also be explained by the competing mutational hazard hypothesis (Lynch et al. 2006), but the evidence for this hypothesis is also very patchy (Smith 2016). Finally, we should not exclude the possibility that our findings may be a statistical artefact caused by the randomness and discontinuity of mitogenomic architecture evolution (Gissi et al. 2008; Zou et al. 2017), as well as the fact that many mitogenomes in public databases are not complete. We attempted to account for the latter problem by conducting the analyses on two different datasets with excluded incomplete mitogenomes and incomplete GOs respectively. Although our conclusions were not affected, we cannot exclude the possibility that the wider size range in endotherms might be an artefact caused by the larger number of available 'endotherm' mitogenomes.

#### Longevity does not have a consistent impact on mitogenomic evolution in flatworms

In further support of this incongruence in evolutionary patterns among different lineages, a study on primates found that generation time (closely associated with longevity) had a negative correlation with GC skew, and positive with the GC content (Min and Hickey 2008). In flatworms, longevity was positively correlated with GC skew magnitude (in disagreement with the working hypothesis), not correlated at all to the GC content, but positively correlated to the GT content (Supplementary figure S1).

#### Parasitic flatworm lineages exhibit higher evolutionary rates than non-parasitic lineages

The prediction of dN/dS values produced noisy results, most probably because this method is designed to test particular hypotheses (relaxation or intensification of selection pressure along a specified set of branches), whereas inferring the exact $\omega$ values for each branch in the dataset using the parameter rich free ratios models is expected to produce unreliable results and the estimates may involve large sampling errors (Yang and Nielsen 2000). When we tested a specific hypothesis using the RELAX tool, we found strong evidence of relaxed purifying selection pressure in parasites. Also, GC skew values corresponded much better to the branch lengths than $\omega$ values (the lowest in non-parasitic species), which further indirectly supports our previous finding that the magnitude of

GC skew is primarily associated with the strength of purifying selection (low absolute skew magnitude = strong purifying selection) (Jakovlić et al. 2021*a*). As absolute $\omega$ values are very difficult to calculate, GC skew magnitude may be a more reliable indicator of the strength of purifying selection pressure, once other factors influencing it are accounted for (Jakovlić et al. 2021*b*).

A factor that should be discussed in more detail is $N_e$, which was proposed as the crucial parameter for explaining the patterns of organellar evolution (Lynch and Conery 2003). Along with the studies cited in the main manuscript, we also hypothesised that parasitic lineages may exhibit reduced $N_e$ in comparison to non-parasitic lineages due to their physical confinement to the body of a single host (Criscione and Blouin 2005; Huyse et al. 2005; Jakovlić et al. 2021*a*), which may cause elevated evolutionary rates. In crustaceans, we found that parasitic species exhibited only nonsignificantly reduced $N_e$ values, and found no evidence for increased speciation rates in parasites (Jakovlić et al. 2021*a*). Indeed, multiple other studies also failed to find support for the central role of $N_e$ in mitogenomic evolution (Bazin et al. 2006; Nabholz et al. 2008; Whitney and Garland 2010). We still cannot reject this hypothesis with full confidence, because our analysis was weakened by the limited availability of molecular data for many lineages, which may have affected both the precision of our $N_e$ estimates and subsequent multilevel regression analyses. Furthermore, $N_e$ estimates can be biased by interlineage variability in the mutation rate, bias towards reflecting recent evolutionary events more than the long-term evolutionary history, and the fact that taxonomic species definition is often problematic in parasites (Daubin and Moran 2004; Whitney and Garland 2010; Hua et al. 2019). Due to these limitations, it is difficult to estimate the impact of $N_e$ with confidence.

It should be mentioned that theoretically mitogenomes of parasites may be affected by selective sweeps as a result of arms races between species (Dawkins and Krebs 1979; Hurst and Jiggins 2005), but this is more likely to affect specific nuclear genes than mitogenomes.
