## Supplementary figures and images for "Drivers of interlineage variability in mitogenomic evolutionary rates in flatworms (Platyhelminthes) are multifactorial"

### Supplementary figure S1

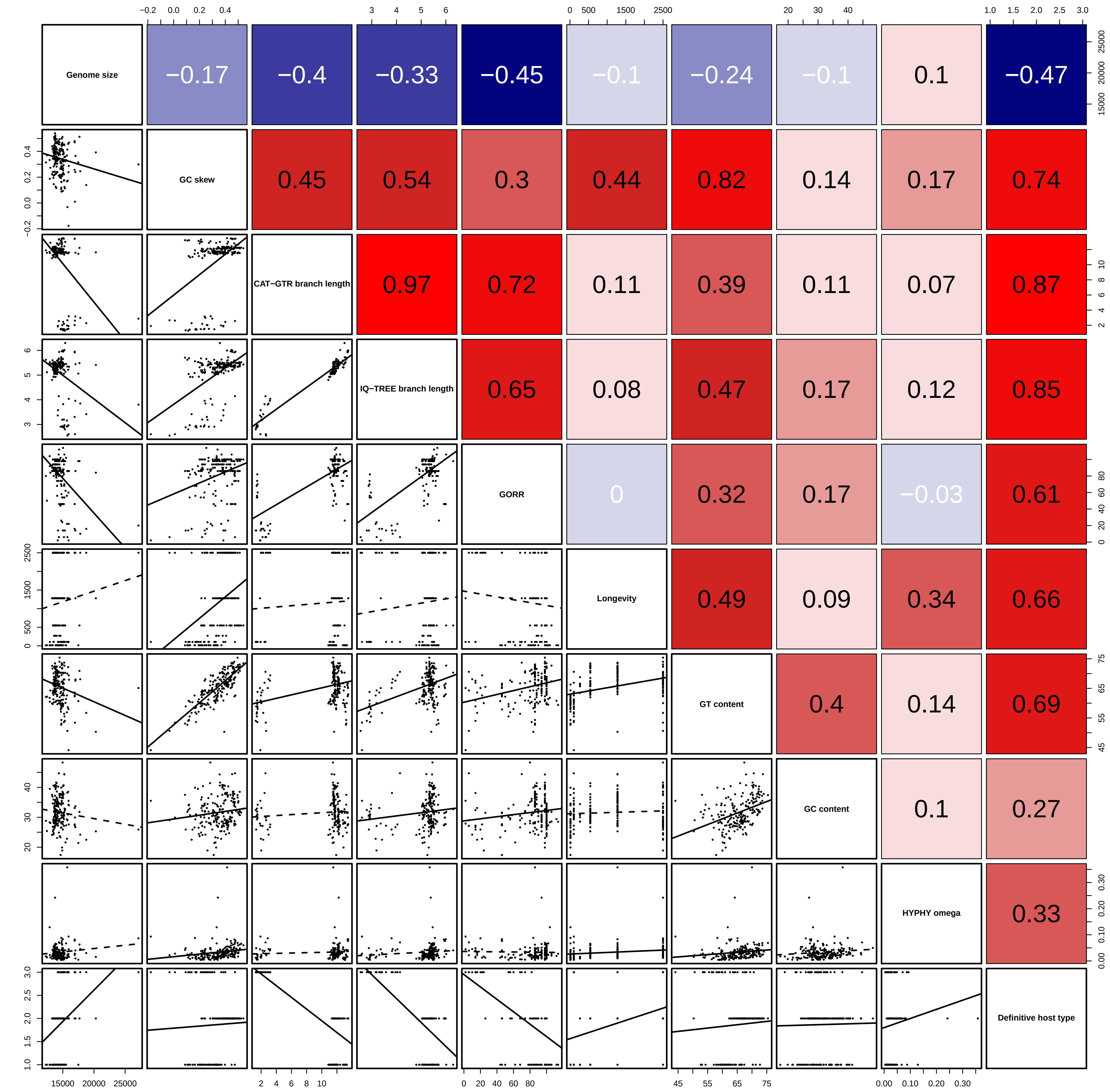

### Supplementary figure S2

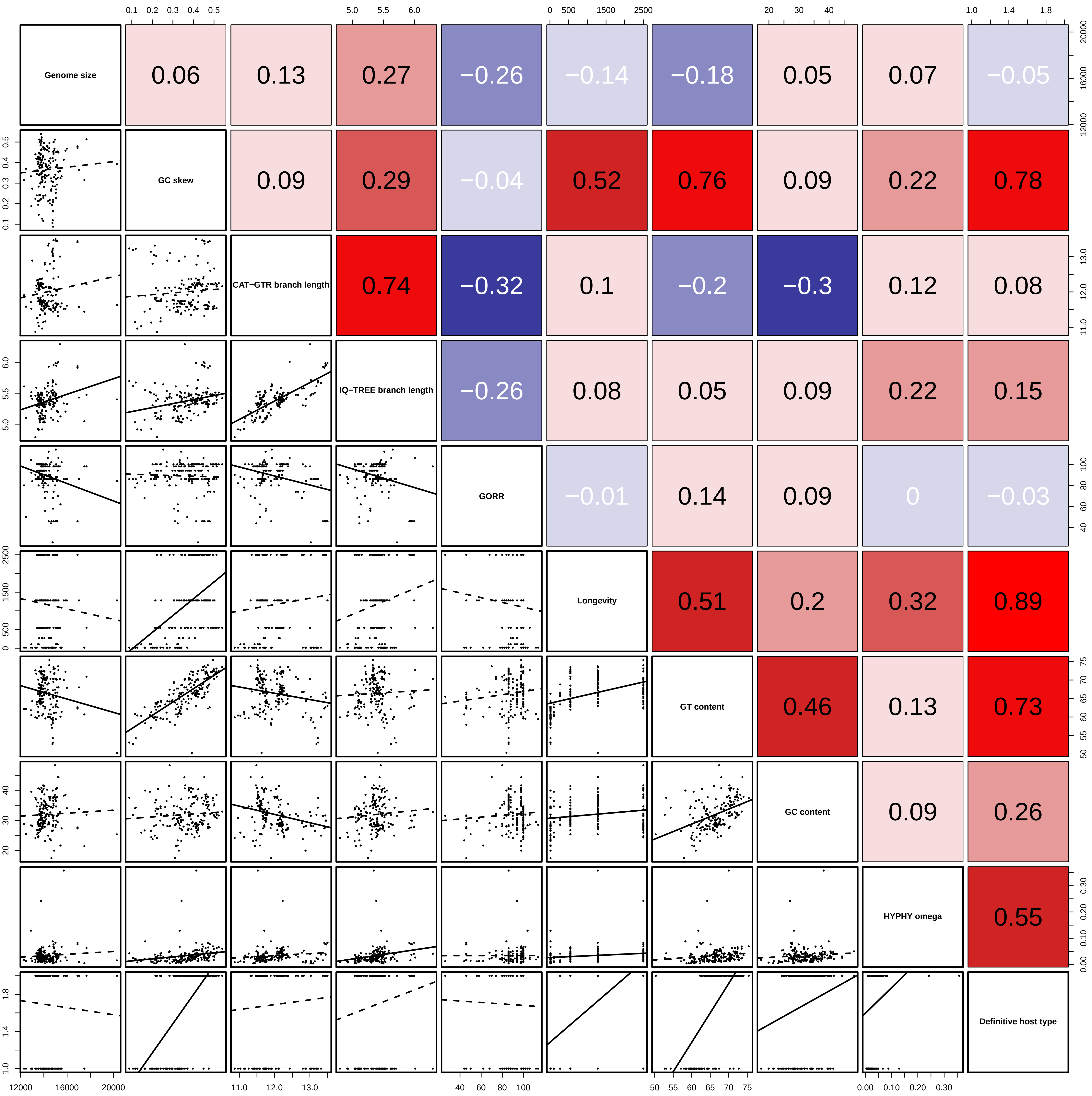

### Supplementary figure S3

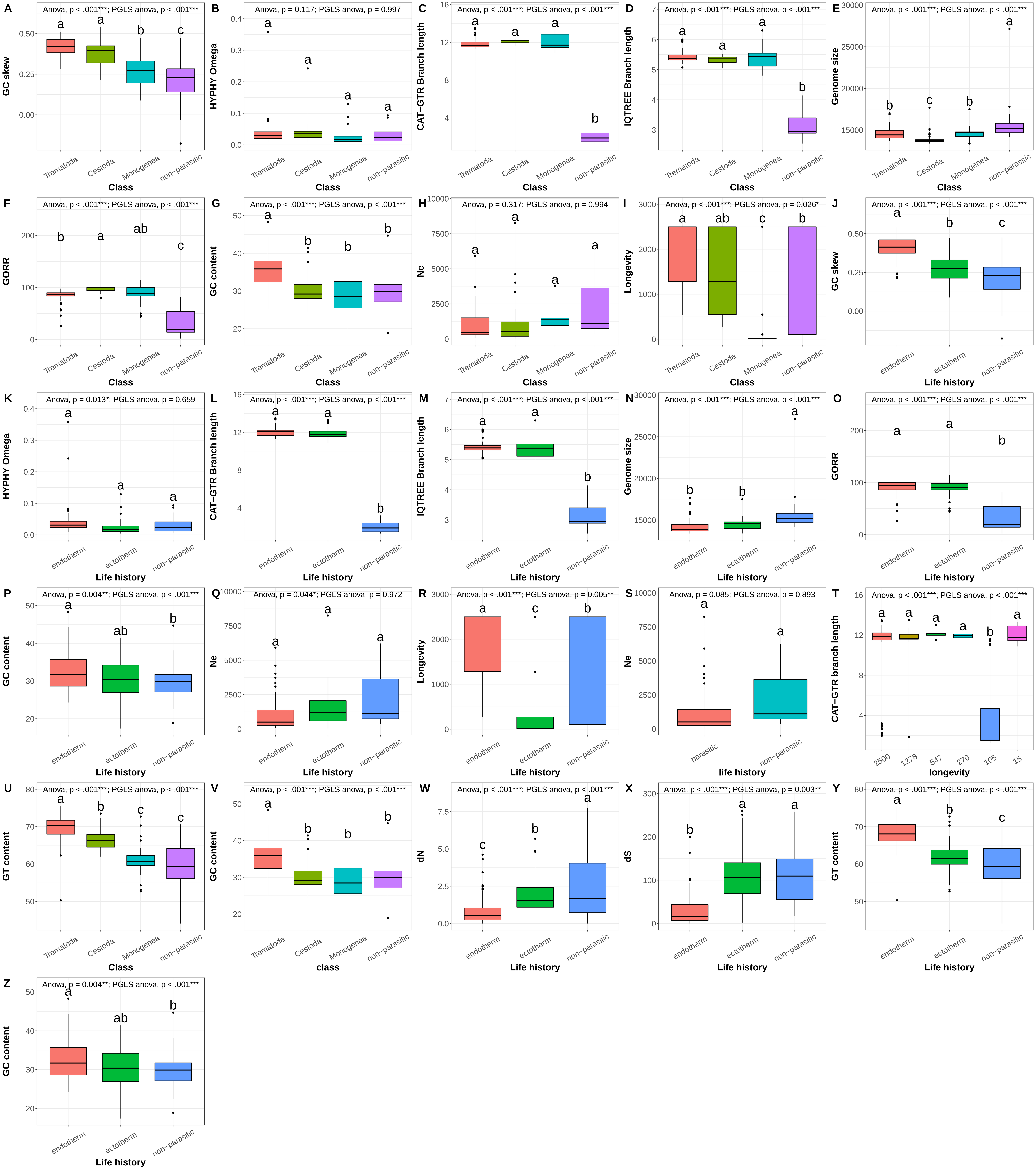
